# Critical role of deadenylation in regulating poly(A) rhythms and circadian gene expression

**DOI:** 10.1101/700443

**Authors:** Xiangyu Yao, Shihoko Kojima, Jing Chen

## Abstract

The mammalian circadian clock is deeply rooted in rhythmic regulation of gene expression. Rhythmic transcriptional control mediated by the circadian transcription factors is thought to be the main driver of mammalian circadian gene expression. However, mounting evidence has demonstrated the importance of rhythmic post-transcriptional controls, and it remains unclear how the transcriptional and post-transcriptional mechanisms collectively control rhythmic gene expression. A recent study discovered rhythmicity in poly(A) tail length in mouse liver and its strong correlation with protein expression rhythms. To understand the role of rhythmic poly(A) regulation in circadian gene expression, we constructed a parsimonious model that depicts rhythmic control imposed upon basic mRNA expression and poly(A) regulation processes, including transcription, deadenylation, polyadenylation, and degradation. The model results reveal the rhythmicity in deadenylation as the strongest contributor to the rhythmicity in poly(A) tail length and the rhythmicity in the abundance of the mRNA subpopulation with long poly(A) tails (a rough proxy for mRNA translatability). In line with this finding, the model further shows that the experimentally observed distinct peak phases in the expression of deadenylases, regardless of other rhythmic controls, can robustly group the rhythmic mRNAs by their peak phases in poly(A) tail length and in abundance of the subpopulation with long poly(A) tails. This provides a potential mechanism to synchronize the phases of target gene expression regulated by the same deadenylases. Our findings highlight the critical role of rhythmic deadenylation in regulating poly(A) rhythms and circadian gene expression.

**Author Summary:** The biological circadian clock regulates various bodily functions such that they anticipate and respond to the day-and-night cycle. To achieve this, the circadian clock controls rhythmic gene expression, and these genes ultimately drive the rhythmicity of downstream biological processes. As a mechanism of driving circadian gene expression, rhythmic transcriptional control has attracted the central focus. However, mounting evidence has also demonstrated the importance of rhythmic post-transcriptional controls. Here we use mathematical modeling to investigate how transcriptional and post-transcriptional rhythms coordinately control rhythmic gene expression. We have particularly focused on rhythmic regulation of the length of poly(A) tail, a nearly universal feature of mRNAs that controls mRNA stability and translation. Our model reveals that the rhythmicity of deadenylation, the process that shortens the poly(A) tail, is the dominant contributor to the rhythmicity in poly(A) tail length and mRNA translatability. Particularly, the phase of deadenylation nearly overrides the other rhythmic processes in controlling the phases of poly(A) tail length and mRNA translatability. Our finding highlights the critical role of rhythmic deadenylation in circadian gene expression control.

## Introduction

Rhythmic control of gene expression is a hallmark of the circadian system. The daily rhythms in biochemistry, physiology and behavior ultimately stem from rhythmic gene expression in each cell [1, 2]. In mammals, approximately 3-15% of mRNAs are rhythmically expressed with a ∼24 hr period in any given tissue [3-5]. The rhythmicity originates from a cell-autonomous circadian clock machinery, which consists of a set of core clock genes interlocked by transcription-translation feedback loops [6-8]. Many core clock genes encode transcription factors and interact with their respective target enhancers to exert rhythmic transcriptional control over circadian mRNA expression [6, 9].

While rhythmic transcriptional control has been extensively studied, rhythmic control of gene expression also occurs beyond transcription [10-12]. Recent genome-wide analyses and mathematical modeling particularly highlight the role of post-transcriptional regulations in driving rhythmic mRNA expression [13-17]. Post-transcriptional regulations target various processes, such as splicing, nuclear export, cellular translocation, dormancy and degradation of RNA [18]. Many post-transcriptional processes are under circadian control; these post-transcriptional processes, in turn, affect the phase and amplitude of mRNA level [10, 19-21].

Ultimately, rhythmic transcription and post-transcriptional processes couple with each other and jointly determine the gene expression rhythm. For example, rhythmic RNA transcription and degradation jointly determine the rhythmicity in mRNA level [16]. As yet, it remains unclear how the rhythmicities in other post-transcriptional processes affect the gene expression rhythm.

One of the post-transcriptional regulations that impact rhythmic gene expression is the regulation of poly(A) tail length. The tracts of adenosines at the 3’ end of nearly all eukaryotic mRNAs are critical for controlling stability and translatability of the mRNAs [22-24]. Hundreds of mRNAs were recently discovered to exhibit robust circadian rhythms in their poly(A) tail lengths in mouse liver [25]. Interestingly, the rhythmicity in poly(A) tail length is closely correlated with the rhythmicity in the corresponding protein level, indicating that rhythmic poly(A) regulation plays an important role in driving rhythmic protein expression [25]. Similar daily fluctuations in poly(A) tail length also occur in mouse brain [26, 27]. In addition, the amplitude of mRNA rhythmicity increases in the absence of *Nocturnin*, a deadenylase (enzyme that removes poly(A) tails from mRNAs) which is rhythmically expressed in different mouse tissues [28, 29]. These data collectively support the importance of poly(A) tail rhythmicity in regulating circadian gene expression.

In this work, we built a mathematical model that describes mRNA dynamics under the regulation of rhythmic transcription, polyadenylation, deadenylation and degradation. We used the model to systematically examine how rhythmic transcription and poly(A) tail regulation generates rhythmicities in poly(A) tail length and mRNA abundance. Our results highlight the rhythmicity in deadenylation as the strongest determinant for the rhythmicities in the poly(A) tail length and in the abundance of mRNAs with long poly(A) tails. The latter can be regarded as a rough proxy for mRNA translatability because the poly(A) tail is known to regulate mRNA translation initiation [30-34]. As a corollary of this general finding, the model further shows that the experimentally observed distinct peak phases in the expression of deadenylases [25] can robustly group the rhythmic mRNAs by their peak phases in poly(A) tail length and in abundance of long-tailed mRNAs, regardless of other rhythmic controls. This result suggests that rhythmic deadenylation can potentially synchronize the rhythmicity in target gene expression regulated by the same deadenylases. Finally, we used the model to predict factors or combination of factors (e.g., amplitudes of or phase differences between specific processes) that can explain the different classes of rhythmic characteristics found in mRNAs with rhythmic poly(A) tail length [25].

## Results

### Model

In a typical RNA expression process, RNAs are first transcribed in the nucleus and acquire long poly(A) tails as a result of nuclear polyadenylation [35]. After being exported into the cytoplasm, mRNAs undergo deadenylation and are ultimately degraded [36]. Cytoplasmic polyadenylation, as another important post-transcriptional regulation, elongates the poly(A) tail to promote mRNA stability and translatability [37]. Although cytoplasmic polyadenylation is typically associated with translational control in oocyte maturation, early embryo development and synaptic plasticity [37-40], it is recently suggested to play a role in circadian gene expression in mouse liver [25]. Furthermore, the expression level of *Gld2*, a poly(A) polymerase responsible for cytoplasmic polyadenylation, exhibits circadian rhythmicity in mouse liver [25]. In light of these biological facts, in the model we incorporated polyadenylation, together with transcription, deadenylation and degradation, to capture the major processes that dynamically regulate poly(A) tail length and mRNA abundance (**Fig. 1A**). Instead of explicitly tracking the exact length of poly(A) tails, the model divides the mRNA population into a long-tailed fraction and a short-tailed fraction, which mimics the experimental condition (long-tailed >∼100nt, short-tailed <∼100nt, [25]). Herein we use the ratio between the abundances of long-tailed and short-tailed mRNAs as the metric for poly(A) tail length, as was done in the experimental study [25].

**Figure 1.**
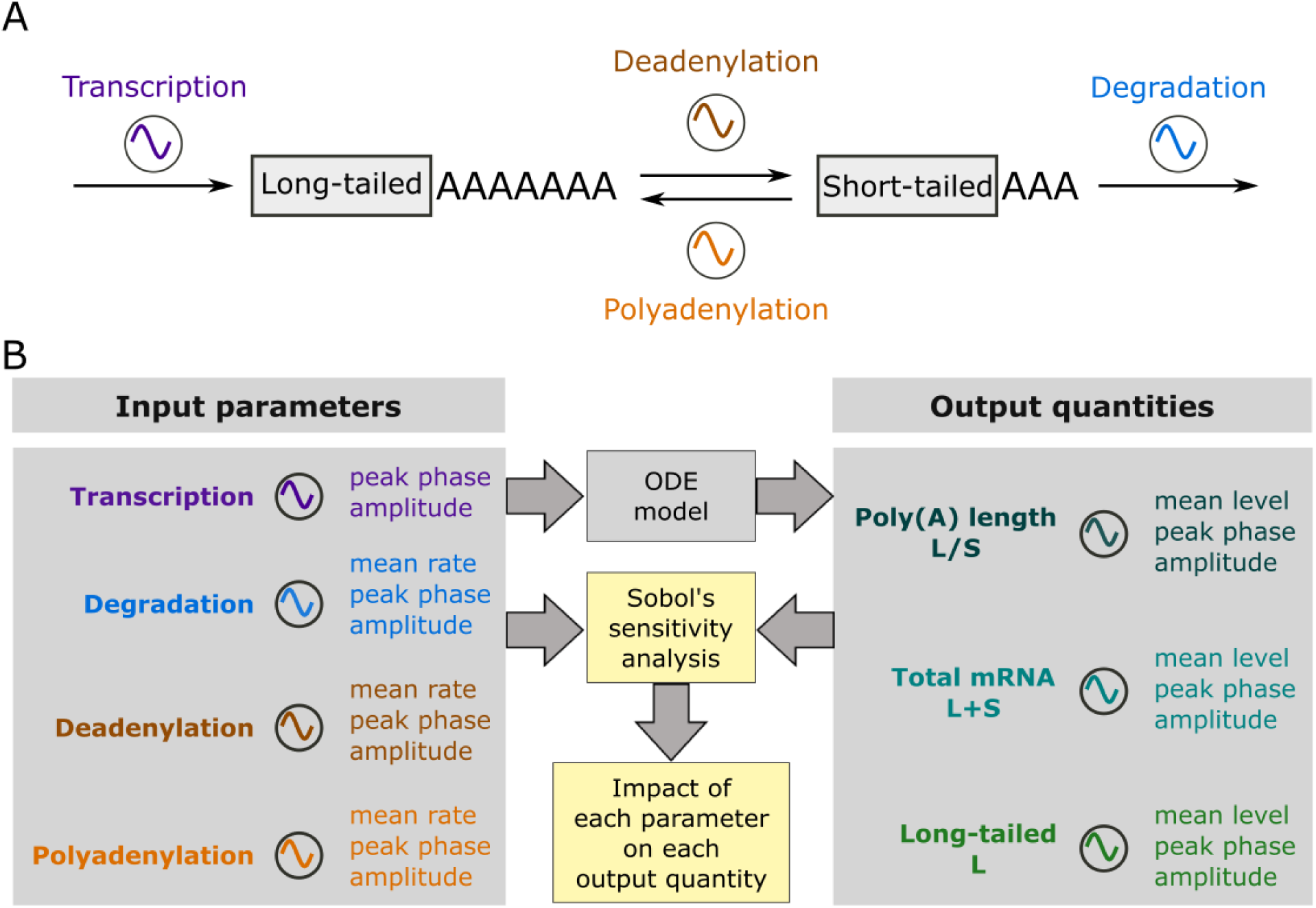
Schematic diagram of the model. (A) The model describes four processes which control the poly(A) tail length and mRNA abundance: transcription, degradation, cytoplasmic deadenylation and polyadenylation. All four processes are allowed variable rhythmicity. (B) Work flow of the study. Numeric simulations of the ODE model using different sets of input parameters (sampled according to **Table 1, Fig. S1**) generate the output quantities. The input parameters and output quantities are analyzed through the global sensitivity analysis to quantify the impact of each parameter on each output quantity over the global parameter space.

For the sake of simplicity, we made the following assumptions in the model based on experimental evidence. First, degradation only occurs to the short-tailed mRNAs, because the poly(A) tail of an mRNA must be shortened to 10∼15 nt before the mRNA is degraded [41-44]. Second, transcription and nuclear polyadenylation are lumped together, because transcription is followed by nuclear polyadenylation in general [45] and the poly(A) polymerases responsible for nuclear polyadenylation are not rhythmically expressed [25]. In our model, therefore, the transcription process directly leads to a long-tailed mRNA, the downstream cytoplasmic deadenylation and polyadenylation further mediate conversion between the long-tailed and short-tailed mRNAs, and degradation consumes the short-tailed mRNA. The ordinary differential equations (ODEs) that govern the temporal dynamics of long-tailed (*L*) and short-tailed (*S*) mRNAs read as Eqs. (1) and (2).

**Table 1.** Parameter distribution for sampling.

| Parameter | Symbol | Distribution | Source |
| --- | --- | --- | --- |
| Mean rate of transcription | $k_{\text{trsc}}$ | 1 (constant) | Has no effect on rhythmic patterns (see Supplementary Materials) |
| Mean rate of degradation | $k_{\text{dgrd}}$ | $\log_{10}(k/\text{hr}^{-1}) \sim \mathcal{N}(-1.10, 0.23^2)$<br>(Log-normal) | Fitting with half-life distribution measured in [56] |
| Mean rate of deadenylation | $k_{\text{deA}}$ | $\log_{10}(k/\text{hr}^{-1}) \sim \mathcal{N}(-0.48, 0.23^2)$<br>(Log-normal) | Mean value of deadenylation rate estimated from [57]; deviation same as mRNA degradation |
| Mean rate of polyadenylation | $k_{\text{polyA}}$ | $\log_{10}(k/\text{hr}^{-1}) \sim \mathcal{N}(-0.48, 0.23^2)$<br>(Log-normal) | Same as deadenylation |
| Relative amplitudes | $A_{\text{trsc}}, A_{\text{dgrd}}, A_{\text{deA}}, A_{\text{polyA}}$ | $A \sim U(0,1)$<br>(Uniform) | |
| Peak phases | $\varphi_{\text{trsc}}, \varphi_{\text{dgrd}}, \varphi_{\text{deA}}, \varphi_{\text{polyA}}$ | $\varphi \sim U(0,24)$<br>(Uniform) | |

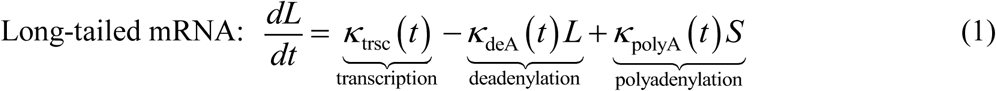

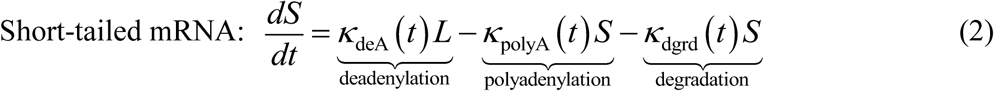

To capture the circadian rhythmicity in the four RNA metabolism processes in Eqs. (1) and (2), each reaction rate term *κ*(*t*) is represented by a sinusoid function like Eq.(3).

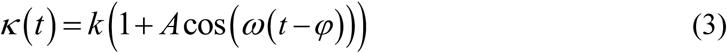

where *k* denotes the mean rate, *A* the relative amplitude, and *φ* the peak phase, of the process labeled by the subscript. The angular frequency, *ω*, equals 2*π/*(24hr). *ω* is fixed, while the other parameters vary. The subscript of a parameter indicates the process it describes (e.g., *k*_deA_ stands for the mean deadenylation rate).

In this work we focus on how rhythmicities in the four processes affect the rhythmicities in total mRNA abundance and poly(A) tail length, because total mRNA abundance and poly(A) tail length were quantified in the previous circadian transcriptome study [13, 25]. Furthermore, we take the rhythmicity of long-tailed mRNA abundance as a rough proxy for mRNA translatability, because poly(A) tail facilitates translation initiation [30-34]. From each simulation result based on Eqs. (1) and (2), i.e., time trajectories *L*(*t*) and *S*(*t*), we extracted the peak phases, relative amplitudes and mean levels of total mRNA abundance (*L*(*t*) + *S*(*t*)), poly(A) length metric (*L*(*t*)*/S*(*t*)), and long-tailed mRNA abundance (*L*(*t*)) (see Methods). These quantities were subject to further analysis, as elaborated in the following Results sections. For the rest of the paper, we will refer to these quantities, e.g., the peak phase of L/S ratio, generally as the “output quantities”, unless any specific quantity is referred to.

### Rhythmic deadenylation is the strongest contributor to rhythmicities in poly(A) tail length and long-tailed mRNA abundance

To investigate the effects of rhythmicities in transcription, degradation, polyadenylation and deadenylation on the output quantities, we ran numeric simulations of the model (Eqs. (1) and (2)) with random parameter values for the mean rates, relative amplitudes, and phases of each process (**Table 1 and Fig. S1**). Only the mean rate of transcription was omitted, because it only affects the overall abundance of mRNAs, but not the output rhythmicity, i.e., phases and relative amplitudes of mRNA abundance and poly(A) tail length (see Supplementary Materials).

Our model results reveal that the peak phase of deadenylation is the strongest contributor to the peak phase of L/S ratio (poly(A) length metric), followed by the peak phase of polyadenylation. Specifically, the scatter plots of the simulation demonstrate a strong correlation between the peak phases of L/S ratio and deadenylation, with a 10 ± 1.5 hr lag between the two (**Fig. 2A**). The peak phase of L/S ratio also depends on the peak phase of polyadenylation, although with much weaker dependence compared to that of deadenylation (**Fig. 2A**). In contrast, the peak phase of L/S ratio depends very little on the peak phases of transcription and degradation (**Fig. 2A**).

**Figure 2.**
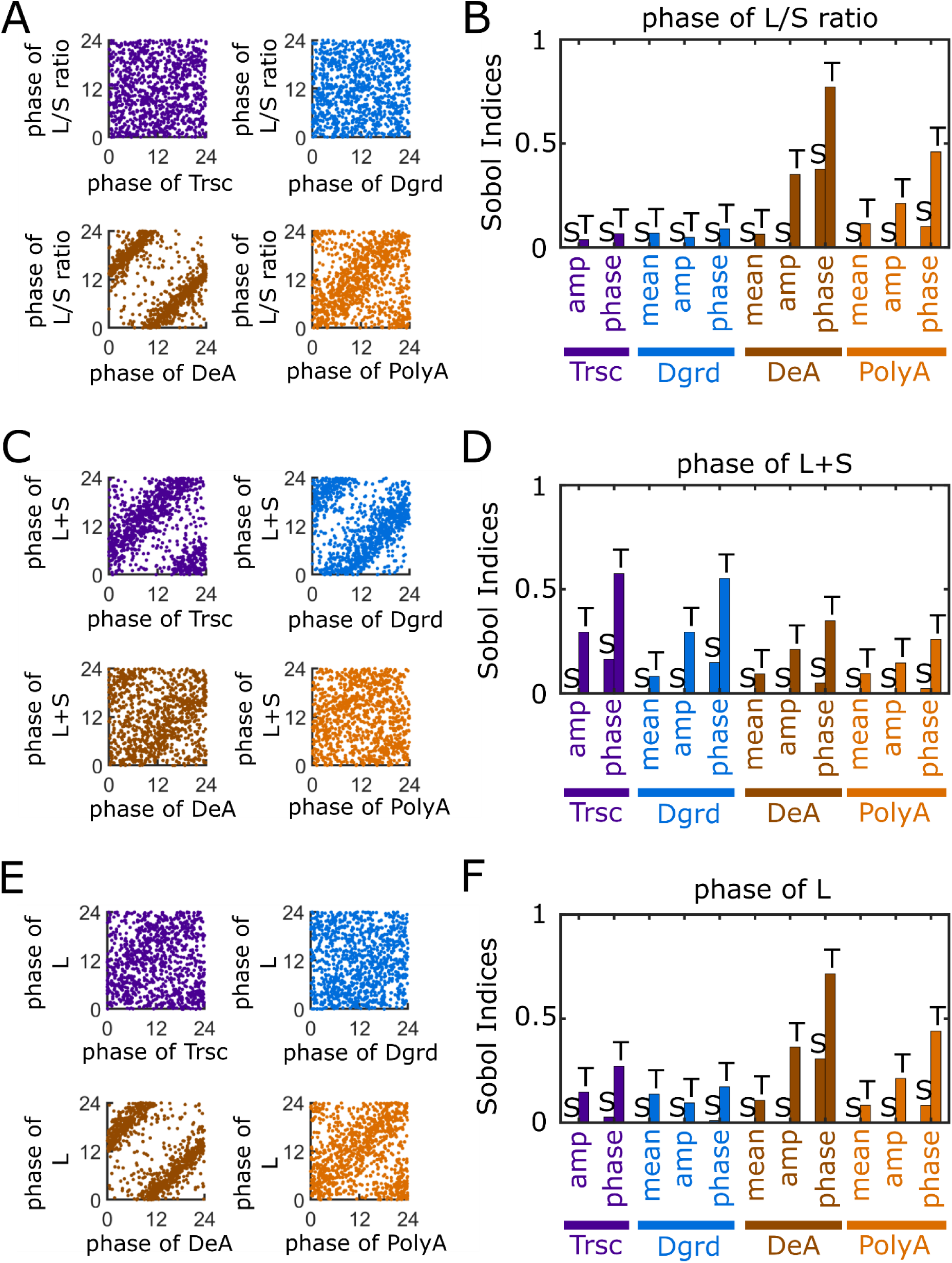
Rhythmicities of poly(A) tail length and long-tailed mRNA abundance are strongly controlled by rhythmic deadenylation. (A) Scatter plot of the peak phases of input processes versus the peak phases of L/S ratio. (B) Sobol indices for the peak phase of L/S ratio (i.e., poly(A) length metric). (C) Scatter plot of the peak phases of input processes versus the peak phases of L+S (i.e., total mRNA abundance). (D) Sobol indices of the peak phase of L+S. (E) Scatter plot of the peak phases of input processes versus the peak phases of L (i.e., long-tailed mRNA abundance). (F) Sobol indices of the peak phase of L. (A, C, E) Each scatter plot shows 10,000 data points randomly chosen from the original simulations for the sake of visual clarity. (B, D, F) Bars with “S” on top: single Sobol indices. Bars with “T” on top: total Sobol indices. Mean values of the Sobol indices are shown, because the variances are too small for clear visualization (**Fig. S2**).

To systematically quantify the impacts of input parameters on output quantities, we performed variance-based sensitivity analysis using the Sobol’s method [46, 47] (**Fig. 1B**, Methods). Based on simulation results from a large number of random parameter sets spanning the global parameter space (**Fig. S1, Table 1**), the Sobol’s method quantifies the sensitivity of an output quantity to an input parameter in terms of how much the parameter, due to the variation in its value, contributes to the variation in the output quantity. Specifically, the sensitivity is reported as the single (S) and total (T) Sobol indices, which represent the contribution of the parameter alone and the contribution of the parameter together with its (nonlinear) interactions with the other parameters, respectively (see Methods).

The estimated Sobol indices (**Fig. 2B**) confirm the findings from the scatter plots (**Fig. 2A**). For example, among all the input parameters, the peak phase of deadenylation has the largest Sobol indices with respect to the peak phase of L/S ratio. The values of the Sobol indices indicate that variance in the peak phase of deadenylation alone contributes to ∼40% of variance in the peak phase of L/S ratio (longest “S” bar in **Fig. 2B**). When the interactions of deadenylation with other processes are counted, this contribution increases to ∼75% (longest “T” bar in **Fig. 2B**). Additionally, the Sobol indices also indicate that the relative amplitude of deadenylation has the strongest impact on the relative amplitude of L/S ratio (**Fig. S2**). In comparison, the mean level of L/S ratio, a quantity not related to rhythmicity, depends nearly equally on the mean rates of deadenylation and polyadenylation (**Fig. S2**). These results collectively demonstrate the rhythmicity in deadenylation as the strongest contributor to the rhythmicity in poly(A) tail length.

Our model results also show a significant impact of rhythmic deadenylation and polyadenylation on the rhythmicity of L+S (total mRNA abundance). Although the peak phases of transcription and degradation strongly influence the peak phase of L+S, as expected (**Figs. 2C, D**), the Sobol indices indicate a weaker, yet substantial impact from the peak phases of deadenylation and polyadenylation on the peak phase of L+S (**Figs. 2D**). These impacts can be understood from the regulation of mRNA stability by poly(A) tail length, which is reflected in the model by the assumption that degradation is restricted to the short-tailed mRNAs (**Fig. 1A**, Eqs. (1) and (2)).

We further used the model to examine the effects of the four processes on the rhythmicity of mRNA translatability, using L (long-tailed mRNA abundance) as a proxy. Although L is a quantity directly related to both L+S level and L/S ratio, the Sobol indices show that the peak phase of L relies most heavily on the peak phase of deadenylation, followed by that of polyadenylation (**Fig. 2F**). Consistently, the scatter plot shows a strong correlation between the peak phases of L and deadenylation, with an approximately 10 hr lag between the two (**Fig. 2E**). This is a relationship highly similar to that observed between the peak phases of L/S ratio and deadenylation (**Fig. 2A**). Furthermore, the relative amplitudes of deadenylation and polyadenylation are also among the strongest contributors to the relative amplitude of L (**Fig. S2**). Overall, the rhythmicities in deadenylation and polyadenylation make stronger impact on the rhythmicity of long-tailed mRNA abundance than the rhythmicities in transcription and degradation. This finding provides an explanation for the observed close correlation between the rhythmicities of poly(A) tail length and protein expression [25].

Cytoplasmic polyadenylation requires specific *cis*-elements in the 3’ untranslated region (UTR) of an mRNA to recruit the molecular machinery that elongates poly(A) tails [38]. However, such *cis*-elements do not necessarily exist in all mRNAs. Therefore, we also removed the polyadenylation term in our model and conducted the same global sensitivity analysis. The results demonstrate similar impacts of the rhythmicity of transcription, deadenylation and degradation on the rhythmicity of L/S ratio, L+S and L (**Fig. S3**) as those found from the model with cytoplasmic polyadenylation (**Figs. 2, 3, S2**). Particularly, rhythmic deadenylation remains the strongest contributor to the rhythmicity of L/S ratio and L. Hence, our conclusion stays the same for mRNAs without the *cis*-elements that mediate cytoplasmic polyadenylation.

**Figure 3.**
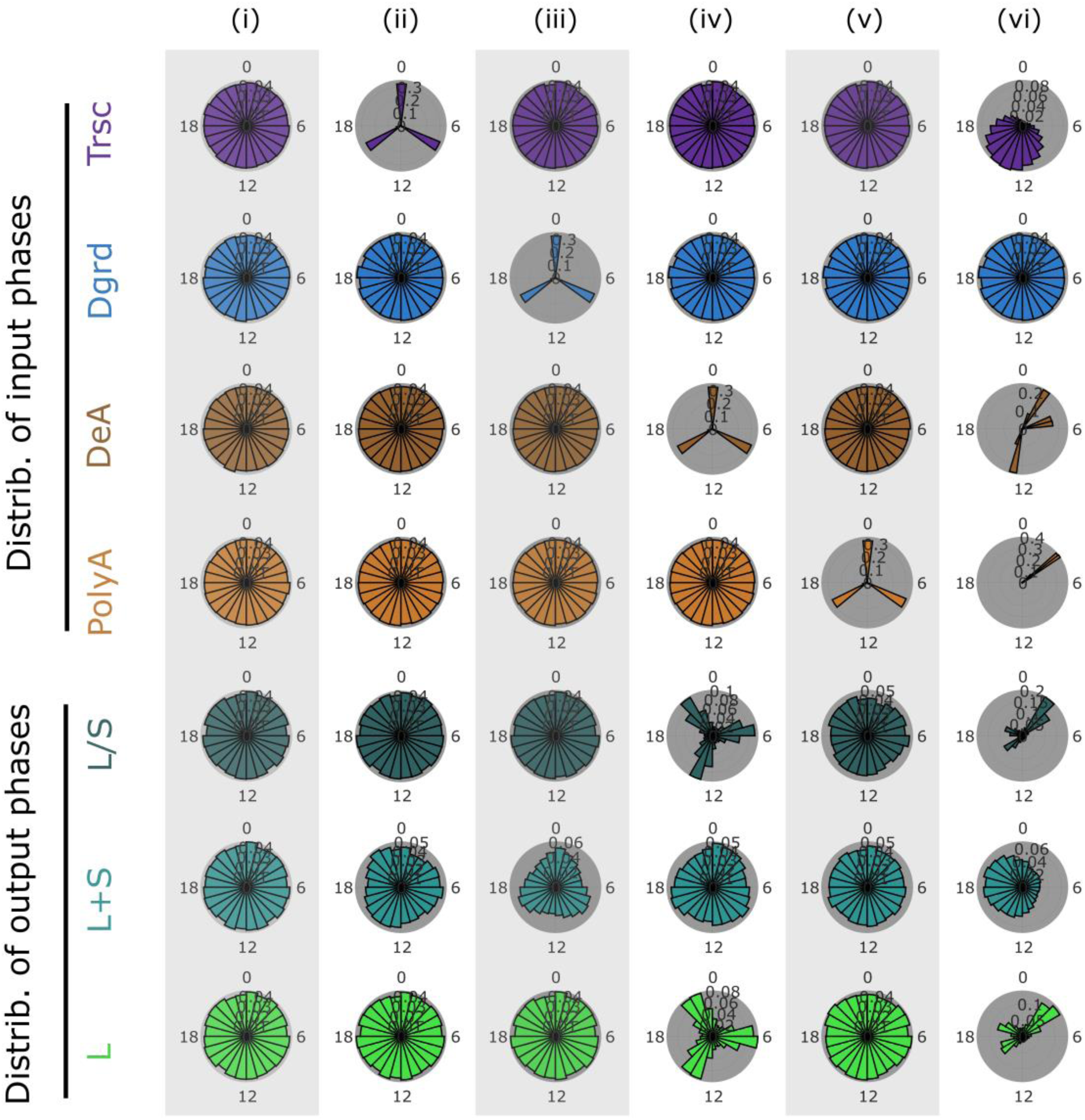
Distinct peak phases in deadenylases group transcripts by their peak phases of poly(A) tail length and long-tailed mRNA abundance. (i) Transcription, degradation, deadenylation and polyadenylation phases evenly distributed around the clock. (ii) Transcription phases within three narrow windows at ZT 0, 8, and 16. Degradation, deadenylation and polyadenylation phases evenly distributed around the clock. (iii) Degradation phases within three narrow windows at ZT 0, 8, and 16. Transcription, deadenylation and polyadenylation phases evenly distributed around the clock. (iv) Deadenylation phases within three narrow windows at ZT 0, 8, and 16. Transcription, degradation and polyadenylation phases evenly distributed around the clock. (v) Polyadenylation phases within three narrow windows at ZT 0, 8, and 16. Transcription, degradation and deadenylation phases evenly distributed around the clock. (vi) Peak phases of transcription follow transcriptome data reported by [13]. Deadenylation phases within three narrow windows at ZT 2, 5, and 13, and polyadenylation phases within one narrow windows at ZT 3.5, based on the data from [25], while degradation phases evenly distributed around the clock. Mean rates and relative amplitudes follow **Table 1** and **Fig. S1**.

Taken together, these model results underscore the importance of rhythmic poly(A) regulation in circadian gene expression, especially its impact on the rhythmicity of poly(A) tail length, total mRNA abundance, and abundance of the long-tailed subpopulation. Importantly, deadenylation emerges as the strongest contributor to the rhythmicity of poly(A) tail length and long-tailed mRNA abundance.

### Rhythmic deadenylation can robustly group genes by their poly(A) tail rhythms

The rhythmicities in transcription, deadenylation, polyadenylation and degradation of mRNAs are ultimately controlled by the rhythmicities in the abundance and activity of the molecules mediating these processes, e.g., transcription factors, deadenylases and poly(A) polymerases. Interestingly, although the core clock machinery includes several transcription factors with different peak phases, the peak phases of nascent RNA synthesis (indicated by intron abundance) are strongly concentrated around ZT 15 in mouse liver [13]. Additionally, a cytoplasmic poly(A) polymerase, *Gld2*, is rhythmically expressed with peak phase around ZT 3.5 (Zeitgeber time, where ZT 0 is defined as the time [hr] of lights on and ZT 12 is defined as the time of lights off) [25]. Meanwhile, five deadenylases are also rhythmically expressed, with *Ccr4e/Angel1* peaking around ZT 2, *Ccr4a/Cnot6, Ccr4b/Cnot6l, Caf1a/Cnot7/pop2* and *Parn* peaking around ZT 5, and *Ccr4c/Nocturnin* peaking around ZT 13 [25]. These data indicate that deadenylases assume a more diverse rhythmic expression pattern than poly(A) polymerases and nascent RNA transcription.

Intrigued by the above observation, we used our model to explore the potential consequence of having several distinct peak phases in deadenylases. In four separate *in silico* experiments, we set transcription, degradation, deadenylation or polyadenylation, respectively, to peak at three narrow windows centered around ZT 0, 8, and 16 (chosen to represent distinct time windows in general), while setting the peak phases of the other three processes to distribute evenly around the clock (**Fig. 3**, ii-v). Our results demonstrate that, when deadenylation peaks in three narrow windows, the peak phases of L/S ratio and L are strongly grouped into three distinct windows as well (**Fig. 3**, iv). In contrast, when transcription (**Fig. 3**, ii), degradation (**Fig. 3**, iii) or polyadenylation (**Fig. 3**, v) peaks in three narrow windows, the resulting peak phases of L/S ratio and L do not show any distinct grouping, and are comparable to the control case with evenly distributed peak phases for all processes (**Fig. 3**, i). To test the effect of the actual rhythmic patterns observed in nascent RNA transcription and expression of deadenylases and polyadenylases, we set the distribution of peak phases centered around ZT 15 for transcription [13], narrow peak phase window centered around ZT 3.5 for polyadenylation, and narrow peak phase windows around ZT 2, ZT 5 and ZT 13 for deadenylation [25]. The simulations results demonstrate that the peak phases of both L/S ratio and L are strongly clustered into three distinct time windows (**Fig. 3**, vi). These results corroborate with the findings above about the strong impact of rhythmic deadenylation on the rhythmicities of L/S ratio and L (**Fig. 2B, F**). Note that the mean rates and relative amplitudes of all four processes assumed random variables in the model simulations (**Table 1, Fig. S1**). Therefore, our results indicate that multiple peak phases in deadenylation, but not other processes, can robustly group the peak phases of poly(A) tail length and mRNA translatability (∼ long-tailed mRNA abundance) into distinct time windows, regardless of variations in the mean rates or rhythmicities of other processes.

### Factors that explain different classes of mRNAs with rhythmic poly(A) tail length

In the previous transcriptome-wide study [25], the mRNAs with <u>p</u>oly(<u>A</u>) tail rhythmicity (PAR mRNAs) were grouped into three classes, based on their rhythmicities in pre-mRNA and total mRNA. The rhythmicity in pre-mRNA essentially reflects the rhythmicity in transcription. The Class I mRNAs are rhythmic not only in poly(A) tail length, but also in pre-mRNA and total mRNA (**Fig. 4A**). The Class II mRNAs are rhythmic in poly(A) tail length and pre-mRNA, but not in total mRNA (**Fig. 4A**). The Class III mRNAs are only rhythmic in poly(A) tail length, but not the other two (**Fig. 5A**). Differences in mRNA half-lives were suggested to explain why the rhythmic patterns of pre-mRNA, total mRNA, and poly(A) tail length are different between these classes [25]. Here we leverage our model to systematically identify factors that can lead to the combinatorial rhythmic patterns in these classes.

**Figure 4.**
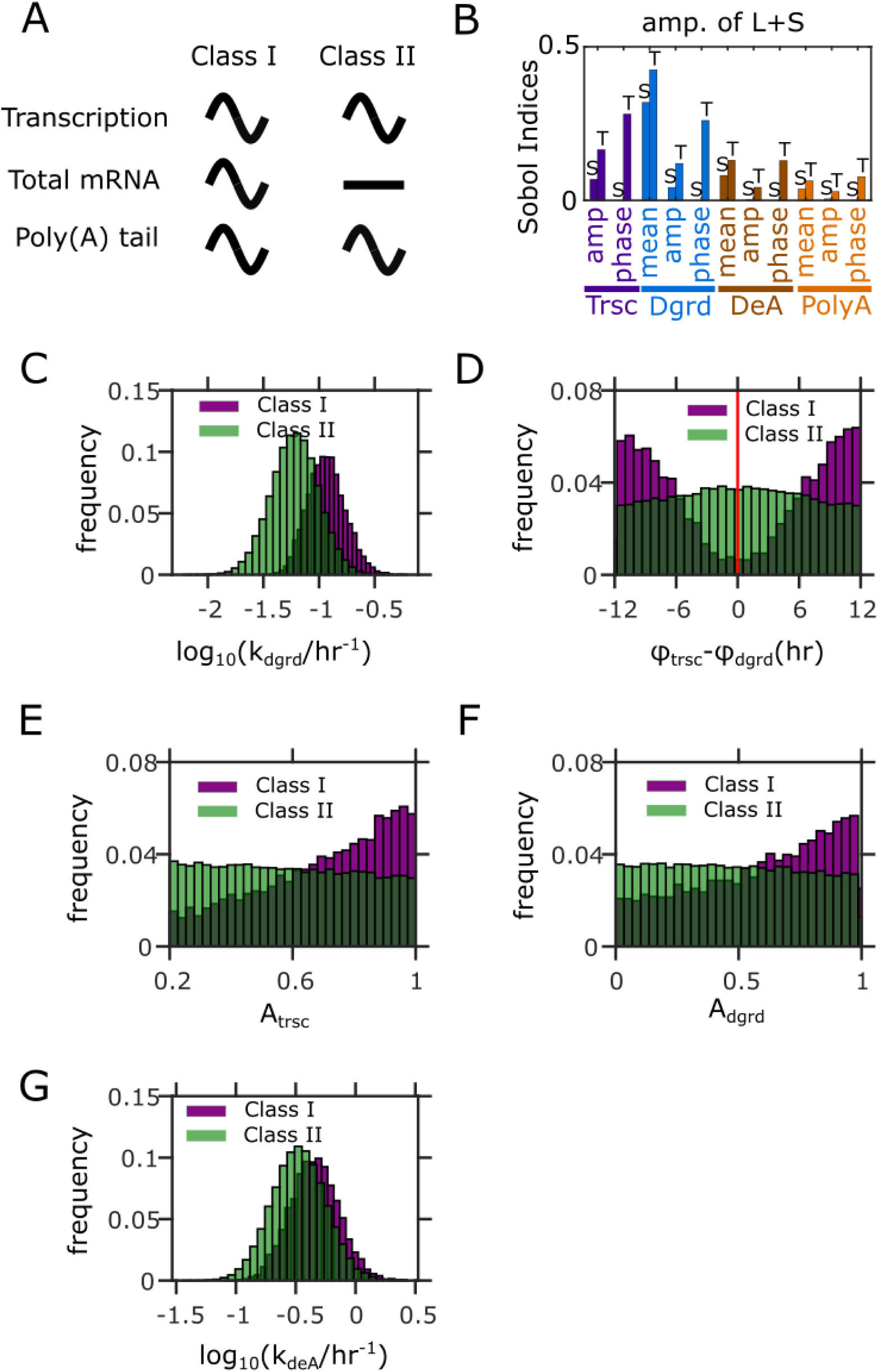
Factors distinguishing between Class I and Class II PAR mRNAs. (A) Characteristics of Class I and Class II PAR mRNAs. (B) Sobol indices for the amplitude of L+S (i.e., total mRNA abundance). Bars with “S” on top: single Sobol indices. Bars with “T” on top: total Sobol indices. (C) Distributions of mean mRNA degradation rates for the two classes. (D) Distributions of peak phase differences between transcription and degradation for the two classes. The red line corresponds to the peak phase of rhythmic degradation. (E) Distributions of relative amplitudes of transcription for the two classes. (F) Distributions of relative amplitudes of degradation for the two classes. (G) Distribution of mean deadenylation rates for the two classes. Results in (C-G) from 100,000 simulations with parameters randomly sampled according to **Table 1**. Parameter sets with *≥ 0.*2 relative amplitude in both L+S and L/S ratio are defined as Class I, while those with < *0.*2 relative amplitude in L+S and *≥ 0.*2 relative amplitude in L/S ratio are defined as Class II.

**Figure 5.**
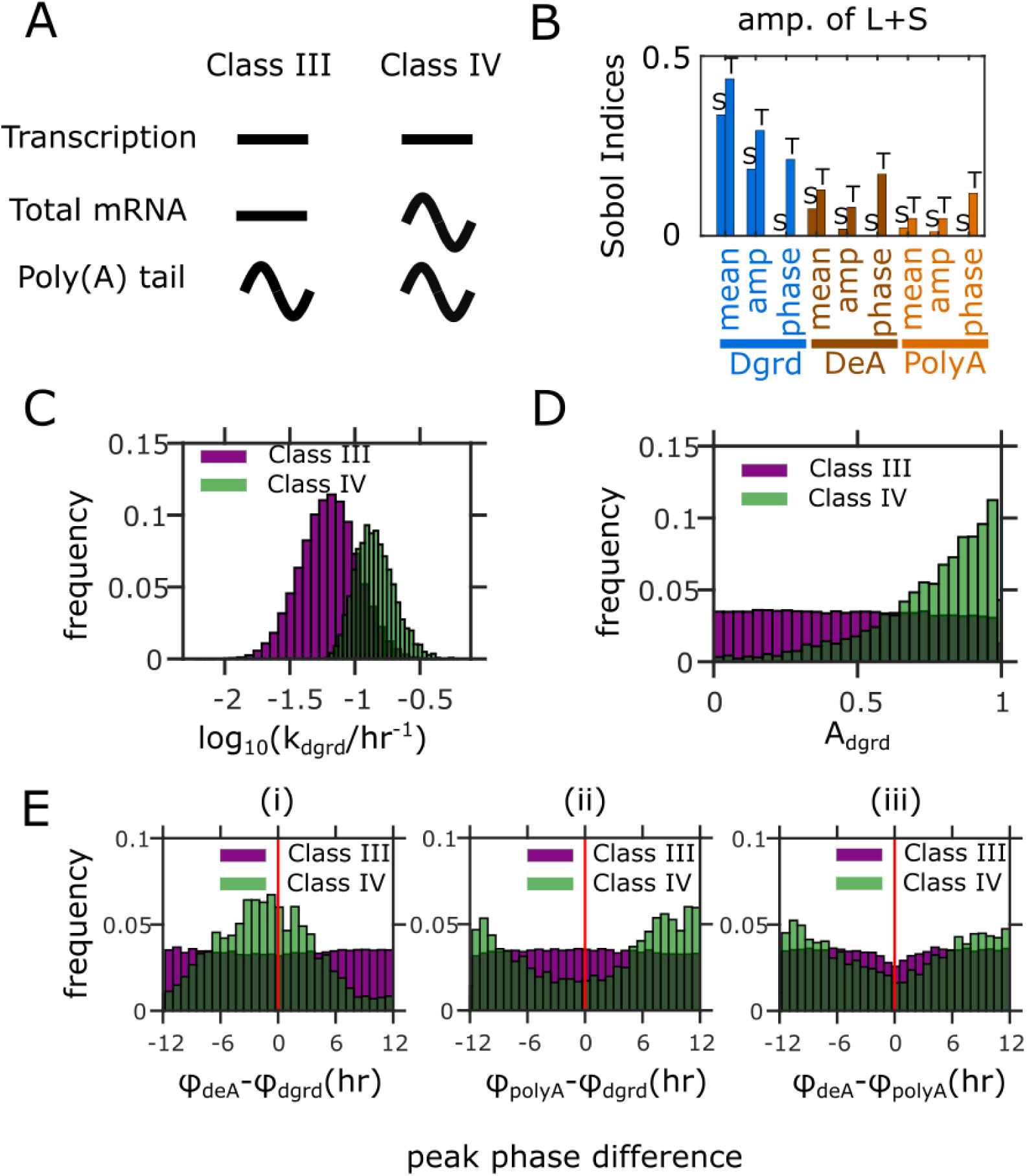
Factors distinguishing between Class III and Class IV PAR mRNAs. (A) Characteristics of Class III and the hypothetical Class IV mRNAs. (B) Sobol indices of the amplitude of L+S (i.e., total mRNA abundance) for the model without rhythmic transcription. Bars with “S” on top: single Sobol indices. Bars with “T” on top: total Sobol indices. (C) Distributions of mean mRNA degradation rates for the two classes. (D) Distributions of relative amplitudes of degradation for the two classes. (E) Distributions of peak phase differences (i) between deadenylation and degradation, (ii) between polyadenylation and degradation, and (iii) between deadenylation and polyadenylation for the two classes. The red lines correspond to the peak phase of degradation (cases (i) and (ii)) or polyadenylation (case (iii)). Results in (C-E) from 100,000 simulations with parameters randomly sampled according to **Table 1**, but without rhythmic transcription (*A*_*trsc*_ = *0*). Parameter sets with *≥ 0.*2 relative amplitude in L/S ratio and < *0.*2 relative amplitude in L+S are defined as Class III, while those with and *≥ 0.*2 relative amplitude in both L/S ratio and L+S are defined as Class IV.

We first attempted to identify the model parameters that contribute most to the distinction between Class I and Class II. Because the only difference between Class I and II is whether total mRNA abundance is rhythmic or not, we focused on identifying model parameters that contribute most to the relative amplitude of L+S. The Sobol indices reveal the mean degradation rate as the strongest contributor to the amplitude of L+S (**Fig. 4B**). We then ran model simulations using random parameter sets (sampled from the distributions given in **Table 1** and **Fig. S1**) and identified the ones that exhibit the characteristics of Class I or Class II (**Fig. 4A**). Out of all the random parameter sets, the mean degradation rates in the Class II parameter sets are overall smaller than those in the Class I parameter sets (**Fig. 4C**). This finding corroborates with the experimental observation that the average half-life (i.e., reciprocal of degradation rate) of Class II mRNAs is longer than that of Class I mRNAs [25].

The total Sobol indices also indicate that the peak phases of transcription and degradation as the second and third strongest contributors to the amplitude of L+S, respectively (**Fig. 4B**). However, the corresponding single indices are diminishingly small (**Fig. 4B**). The huge contrast between the total and single indices indicates that these two parameters exert strong impacts through interactions with other parameters. Because such huge contrasts between total and single indices do not exist in any other parameters, we speculated that the interactions likely happen between the two parameters themselves. Indeed, the Class I, but not the Class II, parameter sets, are strongly enriched with antiphasic rhythms between transcription and degradation (**Fig. 4D**). This finding is consistent with the effect of antiphasic coupling between rhythmic transcription and degradation predicted by a previous modeling study [16].

The Sobol indices also reveal that the relative amplitudes of transcription and degradation rates and the mean deadenylation rate are potentially important contributors to the amplitude of L+S (**Fig. 4B**). Indeed, the Class I parameter sets tend to have stronger amplitudes in transcription and degradation rates (**Fig. 4E, F**), again, consistent with the previous modeling study [16]. Interestingly, unlike the Class I parameter sets (**Figs. 4D-F**, purple), the Class II parameter sets exhibit nearly even distributions of transcription-degradation phase difference, transcription amplitude and degradation amplitude (**Figs. 4D-F**, green). Therefore, generation of significant rhythmicity in L+S (Class I) requires sufficient phase difference and amplitudes in transcription and degradation and sufficiently high amplitudes of transcription and degradation, simultaneously. If any of these conditions are not satisfied, total mRNA abundance would not have significant rhythmicity, and the mRNA would belong to Class II. Lastly, the mean deadenylation rates in the Class I parameter sets tend to be larger than those in the Class II parameter sets (**Fig. 4G**). This is related to the above finding about mRNA half-lives, because deadenylation promotes degradation and hence increasing the mean deadenylation rate has a similar effect on mRNA turnover as increasing the mean degradation rate.

Class III is distinct from Class I and Class II, since it does not have rhythmic transcription (**Fig. 5A**). Because rhythmicity of transcription serves as input to our model, we cannot use the model to identify the origin of lack of transcriptional rhythmicity. However, we are interested in understanding why all PAR mRNAs without transcriptional rhythmicity also lack rhythmicity in L+S, and only exhibit rhythmicity in L/S ratio [25]. For the convenience of discussion, we use “Class IV” to refer to a hypothetical group of PAR mRNAs that would exhibit rhythmicity in total mRNAs and poly(A) tails, but not in pre-RNA (**Fig. 5A**); this group of mRNAs are not found in the experiments [25]. We used the model to identify model parameters that could contribute to the difference between Class III and the hypothetical Class IV. Because both Class III and Class IV do not have rhythmic transcription, we ran model simulations with non-rhythmic transcription (i.e., setting the relative amplitude of transcription to zero, while keeping the other parameters sampled from the same distributions as before (**Table 1, Fig. S1**)). Out of the random parameter sets, we identified those that fit the characteristics of Class III or Class IV (**Fig. 5A**). We also calculated the Sobol indices for this model.

When the model does not have rhythmic transcription, the Sobol indices again reveal the mean degradation rate as the strongest contributor to the relative amplitude of L+S (**Fig. 5B**). Consistently, the Class IV parameter sets require much larger mRNA degradation rate, i.e., much shorter half-life of mRNA, than the Class III parameter sets, to sustain rhythmic total mRNA (**Fig. 5C**). Therefore, the absence of Class IV mRNAs in the experiment is most likely due to the long half-lives of the mRNAs without rhythmic transcription. Indeed, Class III has the longest average mRNA half-life measured among all mRNAs that are rhythmically expressed [25].

We also identified a few additional factors that would distinguish Class III from Class IV. The second strongest factor affecting the amplitude of L+S is the relative amplitude of degradation rate, based on the Sobol indices (**Fig. 5B)**. The Class IV parameter sets have markedly higher amplitudes of degradation rate than Class III (**Fig. 5D**). The phases of all three rhythmic processes, i.e., degradation, deadenylation and polyadenylation, are also potentially important contributors, because their total Sobol indices are substantial (**Fig. 5B**). Again, the huge contrast between the total and single indices for these phase parameters, but not the other parameters, suggests that they exert impacts through interactions among themselves. We hence examined the distribution of pairwise differences between the three phase parameters. To achieve the rhythmic characteristics of the hypothetical Class IV, the peak phases of deadenylation and degradation need to be close to each other, but opposite to that of polyadenylation (**Fig. 5E**). This can be understood from the fact that both deadenylation and degradation promote mRNA turnover while polyadenylation inhibits it. Unlike the Class IV parameter sets, no distinct patterns are found in the amplitude of degradation rate or the phase differences in the Class III parameter sets (**Figs. 5D, E**). Similar to the discussion above for Class I and Class II, these results indicate that the Class IV characteristics requires both sufficiently large amplitude in degradation and sufficient differences of the polyadenylation phase from the deadenylation and degradation phases. The missing of Class IV from the experiment suggests that mRNAs without transcriptional rhythmicity do not satisfy these conditions at the same time.

Overall, our model suggests that besides mRNA half-life, relative amplitudes and phase difference between transcriptional and post-transcriptional processes also contribute to the rhythmic characteristics that distinguishes the three observed classes of PAR mRNAs (**Figs. 4, 5**). These results highlight that rhythmic transcriptional and post-transcriptional processes collectively determine the rhythmicity in mRNA expression and poly(A) tail length. It will be of future interests to test if the factors predicted by the model are indeed correlated with different rhythmic characteristics.

## Discussion

Many post-transcriptional steps are involved in regulating circadian gene expression [10], among which rhythmic regulation of poly(A) tail length was recently discovered and characterized at the genome-wide scale [25]. Notably, the rhythmicity in poly(A) tail length can be both the cause and result of rhythmic gene expression. On the one hand, the rhythmicity in poly(A) tail length can control the rhythmicity in mRNA degradation [16], because mRNA stability is related to the poly(A) tail length [24, 30-32]. On the other hand, the rhythmicity in poly(A) tail length itself stems from the rhythmic control of post-transcriptional processes, such as polyadenylation and deadenylation. At the mechanistic basis, it is the dynamic coupling between multiple rhythmic processes, i.e., transcription, polyadenylation, deadenylation and degradation, that determines the rhythmic patterns in both poly(A) tail length and mRNA abundance.

In this work, we developed a parsimonious mathematical model (**Fig. 1**) to quantitatively evaluate how rhythmic inputs from the transcription, degradation, polyadenylation and deadenylation processes collectively determine the rhythmic outputs in mRNA abundance, poly(A) tail length and long-tailed mRNA abundance. Our model results and global sensitivity analyses reveal rhythmic deadenylation as the strongest factor in controlling the peak phases and amplitudes of rhythmic poly(A) tail length and long-tailed mRNA abundance (**Figs. 2, 3**). This finding highlights the crucial role of rhythmic poly(A) regulation in circadian gene expression. Our model also suggests how three classes of rhythmic characteristics observed in PAR mRNAs [25] arise from the dynamic features of the four processes, as well as the coupling among their rhythmicities (**Figs. 4, 5**).

The importance of dynamic coupling between rhythmic transcription and post-transcriptional processes was demonstrated by a previous modeling study by Lück et al. [16]. That work particularly highlights that rhythmic turnover is necessary for achieving >6 hr peak phase difference between transcription and mRNA abundance. In comparison, our study explicitly considers the effects of poly(A) regulation, a common intermediate process in the mRNA decay pathway, on the rhythmicity of both mRNA abundance and poly(A) tail length. Because deadenylation is necessary for mRNA degradation and polyadenylation opposes it, rhythmic deadenylation and polyadenylation, unsurprisingly, affect the rhythmicity of total mRNA abundance at a level comparable to rhythmic degradation (**Figs. 2, S2**). However, when poly(A) tail length and its effect on mRNA translatability are considered, rhythmic deadenylation emerges as the most important step affecting their rhythmicity (**Figs. 2, S2**). Of course, our model has not included other mRNA decay pathways that do not depend on poly(A) regulation, such as endonuclease cleavage of mRNA followed by 5’-3’ decay [42]. For any mRNA decayed through these pathways, which are less common, their rhythmicity obviously would not depend on the rhythmicity in poly(A) regulation.

Based on the finding of rhythmic deadenylation as the strongest contributor to rhythmicity of poly(A) tail length and long-tailed mRNA abundance, we further discovered that rhythmic deadenylation is capable of synchronizing the target circadian gene expression post-transcriptionally. According to the model results, three distinct peak phases in deadenylation activity, as those observed in mouse liver [25], can robustly divide the mRNAs into three distinct groups by their peak phases of poly(A) tail length and long-tailed mRNA abundance; this grouping effect by deadenylation phases happens regardless of the rhythmicity in other processes (**Fig. 3**). This finding suggests a potential mechanism to synchronize the expression of genes controlled by the same deadenylases and hence foster synergy among these genes around the clock. This synchronization potential is unique to rhythmic deadenylation, but not the other rhythmic processes (**Fig. 3**).

The capability of deadenylation to synchronize circadian gene expression further poses two interesting questions. First, could deadenylation help synchronize circadian gene expression among different cells and entrain the cell-autonomous clock to the systemic rhythms? Recent studies suggest that rhythmic feeding or other systemic rhythmic cues control the rhythmic expression of several deadenylases, including *Parn, Pan2* [17] and *Nocturnin* [48], through clock-independent pathways. Given the findings by our model, such systemically driven rhythmicity in deadenylases could dictate the rhythmicity of poly(A) length and mRNA translatability (∼long-tailed mRNA abundance). This could help synchronize circadian gene expression in cells influenced by the same systemic signals. Second, could deadenylases play a role in tissue-specific circadian gene expression? Rhythmic gene expression is known to vary tremendously from tissue to tissue: different tissues not only share very few rhythmically expressed genes beyond the core clock genes, but also display different peak times for some genes [5, 49, 50]. It is puzzling how the rhythmicity in gene expression varies so much across different tissues while the cellular clock machineries are the same and are presumably synchronized throughout the organism. Most previous studies on the mechanisms of tissue-specific circadian gene expression have focused on tissue-specific transcriptional control, such as rhythmic fluctuations in chromatin structure and interactions between core clock transcription factors and tissue-specific transcription factors [51, 52]. In light of the findings from our work, differential expression patterns of deadenylases in different tissues [53] could serve as an additional mechanism to mediate tissue-specific circadian gene expression. These two interesting questions await future studies to answer.

Circadian gene expression is a critical, yet highly complex process. Expressing the right gene at the right time and the right place requires coordinated control at various gene expression steps, as well as across different cells and tissues. Systems-level study of the coupling between different rhythmic processes is necessary to gain comprehensive understanding of circadian gene expression control, and more importantly, the ability to make positive use of circadian rhythm in treatments of diseases. As our work demonstrates the significant impact of rhythmic poly(A) regulation and its coupling with rhythmic mRNA transcription and degradation on circadian gene expression, it will be of great future interest to examine how coupling between the rhythmicities of transcription, poly(A) regulation and other post-transcriptional, translational and post-translational processes influences circadian gene expression.

## Methods

### Model simulation and extraction of phase, amplitude and mean from simulation results

For any given parameter set, Eqs. (1) and (2) were simulated using the ordinary differential equation (ODE) solver, ode45, in MATLAB. For a simulated time trajectory {*L*(*t*), *S*(*t*)}, the peak phases, relative amplitudes and mean levels of *L*(*t*) + *S*(*t*), *L*(*t*)*/S*(*t*) and *L*(*t*) were analyzed. First, the time trajectory for the output quantity of interest, e.g., *L*(*t*)*/S*(*t*), was calculated from {*L*(*t*), *S*(*t*)}. Then a 48-hr window after 700 hrs (sufficiently long to pass the initial transient) were extracted from the simulated trajectory for data analysis. The trajectories typically have irregular time spacing (due to automatic time stepping in the ode45 solver) and often have insufficient time resolution for accurate determination of the peak phase. To improve the accuracy of the estimated peak phase, the 48-hr trajectory were interpolated upon 500 equally spaced time points spanning the 48 hrs. The peak phase was evaluated from the time for the maximum value, *t*_*max*_, i.e., peak phase = *mod*(*t*_*max*_ + 700, 24) (hr). The mean value was estimated by taking the average of the 500 interpolated data values within the 48-hr window. The relative amplitude was estimated by taking the maximum and minimum values within the 48-hr window and calculating (max − min)*/*(2 × mean). An output quantity was considered rhythmic if its relative amplitude is equal to or greater than 0.2.

### Parameter sampling

We performed global parameter sensitivity analysis [54] on the model to analyze the general contribution of each parameter to any specific output quantity (i.e., peak phase, relative amplitude and mean of *L*(*t*) + *S*(*t*), *L*(*t*)*/S*(*t*) and *L*(*t*)). To perform such global sensitivity analysis, one needs to simulate the model with randomly chosen parameter values that maximally represent the parameter space. In this study we drew random parameter values from the distributions listed in **Table 1** and plotted in **Fig. S1**. The peak phases and relative amplitudes were sampled form uniform distributions of their possible ranges by definition (**Table 1**). The mean reaction rates were sampled from log-normal distributions suggested by previous genomic scale measurements (see sources indicated in **Table 1**). It is noteworthy that we set the mean transcription rate as constant, as it only causes proportional changes in *L*(*t*) and *S*(*t*), and does not affect the rhythmic patterns of any quantity (see Supplementary Materials). To improve the accuracy of the global sensitivity analysis for models with many parameters, one needs sampled parameter values that well represent the parameter space. To this end, we used the sampling method of Latin hypercube [55], which is known to ensure good representation of the high-dimensional parameter space.

### Sobol’s method of global sensitivity analysis

To evaluate the impact of each model parameter (e.g., phase of deadenylation) on each model output (e.g., relative amplitude of L/S ratio), we used a variance-based global parameter sensitivity analysis method, the Sobol indices [47]. For each pair of parameter *X*_*i*_ and output *Y*_*j*_ in the model, the first-order, or single Sobol index, *S*_*ij*_, characterizes the contribution of variance in *X*_*i*_ alone to the total variance in *Y*_*j*_ (Eq.(4)). The total-effect, or total index, *S*_*Tij*_, characterizes the contribution of variance in *X*_*i*_, as well as the variance caused by its coupling with other parameters, to the total variance in *Y*_*j*_ (Eq.(5)). The larger these indices are, the more sensitive *Y*_*j*_ is to *X*_*i*_, or the more impact *X*_*i*_ has on *Y*_*j*_.

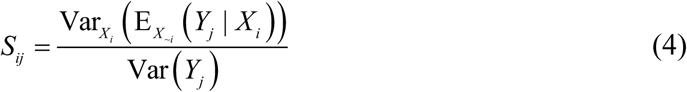

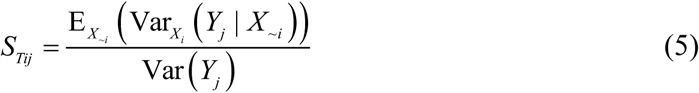

where the subscript ∼*i* indicates all indices except for *i*.

We followed the specific algorithms given in [46] and [58] for evaluating the single (Eq.(4)) and total indices (Eq.(5)), respectively. The details of implementation are explained below.

1. Sample from the distributions given in **Table 1** two independent groups of *N* parameter sets (*N* = 100,000 in this study):

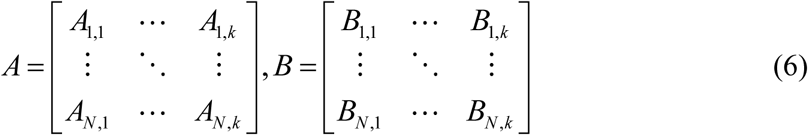

Each row in *A* and *B* represents one set of *k* parameters. *k* = 11 for the model with cytoplasmic polyadenylation. *k* = 8 for the model without cytoplasmic polyadenylation. *k* = 9 for the model without transcriptional rhythmicity.
2. Construct *k* hybrid groups of parameter sets. The *i*-th group, 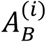, has the *i*-th column equal to the *i*-th column of *B*, and the remaining columns copied from *A*, where *i* = 1, …, *k*.

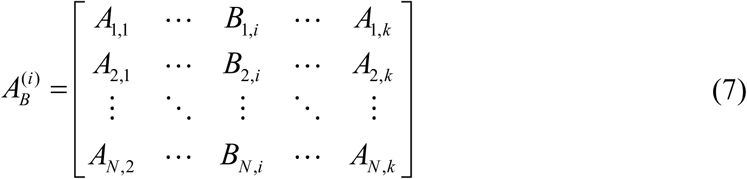
3. Estimate the total variance in the output.

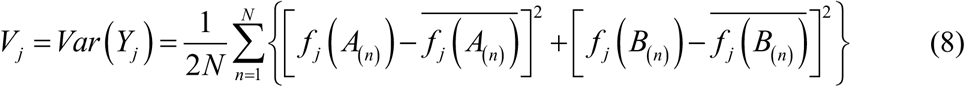

where *f*_*j*_ denotes the *j*-th output quantity from the circadian gene expression model (Eqs. (1) and (2)). *A*_(*n*)_ and *B*_(*n*)_ denote the *n*-th parameter set (row) in Groups *A* and *B*, respectively. The bars on top denote the average of output quantities over *N* parameter sets.
4. Estimate the single and total Sobol indices, using Eqs. (9) and (10) [46, 58].

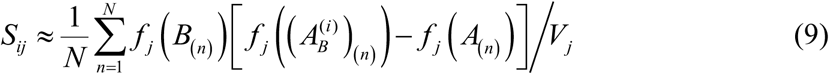

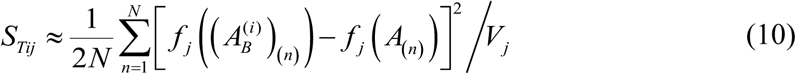

where 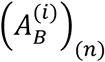 denotes the *n*-th parameter set (row) in the *i*-th hybrid group, and the other notations follow those described above.

## Acknowledgement

We thank Dr. Xi Chen (Virginia Tech) for helpful discussion of the Sobol method.

## Supporting Information

**S1 File: Supplementary Materials.**

