## Supplementary Materials for "Critical role of deadenylation in regulating poly(A) rhythms and circadian gene expression"

**Setting mean transcription rate as constant does not affect rhythmic pattern**

Eqs. (1) and (2) from the Methods section are copied below.

Long-tailed mRNA: $$\frac{dL}{dt} = \kappa_{\text{trsc}}(t) - \kappa_{\text{deA}}(t)L + \kappa_{\text{polyA}}(t)S \tag{S1}$$

Short-tailed mRNA: $$\frac{dS}{dt} = \kappa_{\text{deA}}(t)L - \kappa_{\text{polyA}}(t)S - \kappa_{\text{dgrd}}(t)S \tag{S2}$$

Note that $\kappa_{\text{trsc}}(t) = k_{\text{trsc}}\,(1 + A_{\text{trsc}}\cos(\omega(t - \varphi_{\text{trsc}})))$. We can divide both sides of Eqs. (S1) and (S2) by the mean transcription rate constant, $k_{\text{trsc}}$ and obtain the following equations:

$$\frac{d\left(L/k_{\text{trsc}}\right)}{dt} = \left(1 + A_{\text{trsc}}\cos\left(\omega t - \varphi_{\text{trsc}}\right)\right) - \kappa_{\text{deA}}(t)\frac{L}{k_{\text{trsc}}} + \kappa_{\text{polyA}}(t)\frac{S}{k_{\text{trsc}}} \tag{S3}$$

$$\frac{d\left(S/k_{\text{trsc}}\right)}{dt} = \kappa_{\text{deA}}(t)\frac{L}{k_{\text{trsc}}} - \kappa_{\text{polyA}}(t)\frac{S}{k_{\text{trsc}}} - \kappa_{\text{dgrd}}(t)\frac{S}{k_{\text{trsc}}} \tag{S4}$$

Eqs. (S3) and (S4) show that changing $k_{\text{trsc}}$ only causes a strictly proportional change of $L(t)$ and $S(t)$. The rhythmicity patterns, including the peak phase and relative amplitude, will not be affected at all. When $k_{\text{trsc}}$ changes, the means of $L(t)$ and $S(t)$ change proportionally, but mean L/S ratio remains the same.

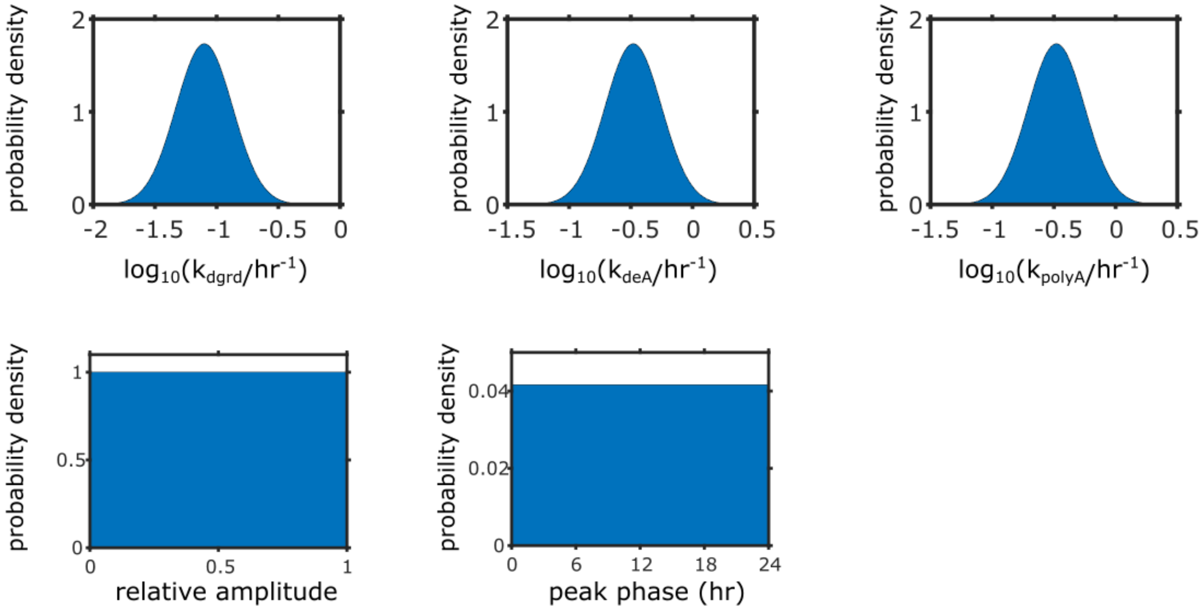


**Figure S1. Sampling distribution of the parameters in model.** (A) Sampling distribution of mean mRNA degradation rates. (B) Sampling distribution of mean deadenylation rates. (C) Sampling distribution of mean polyadenylation rates. (D) Sampling distribution of relative amplitudes of all rhythmic processes. (E) Sampling distribution of peak phases of all rhythmic processes.

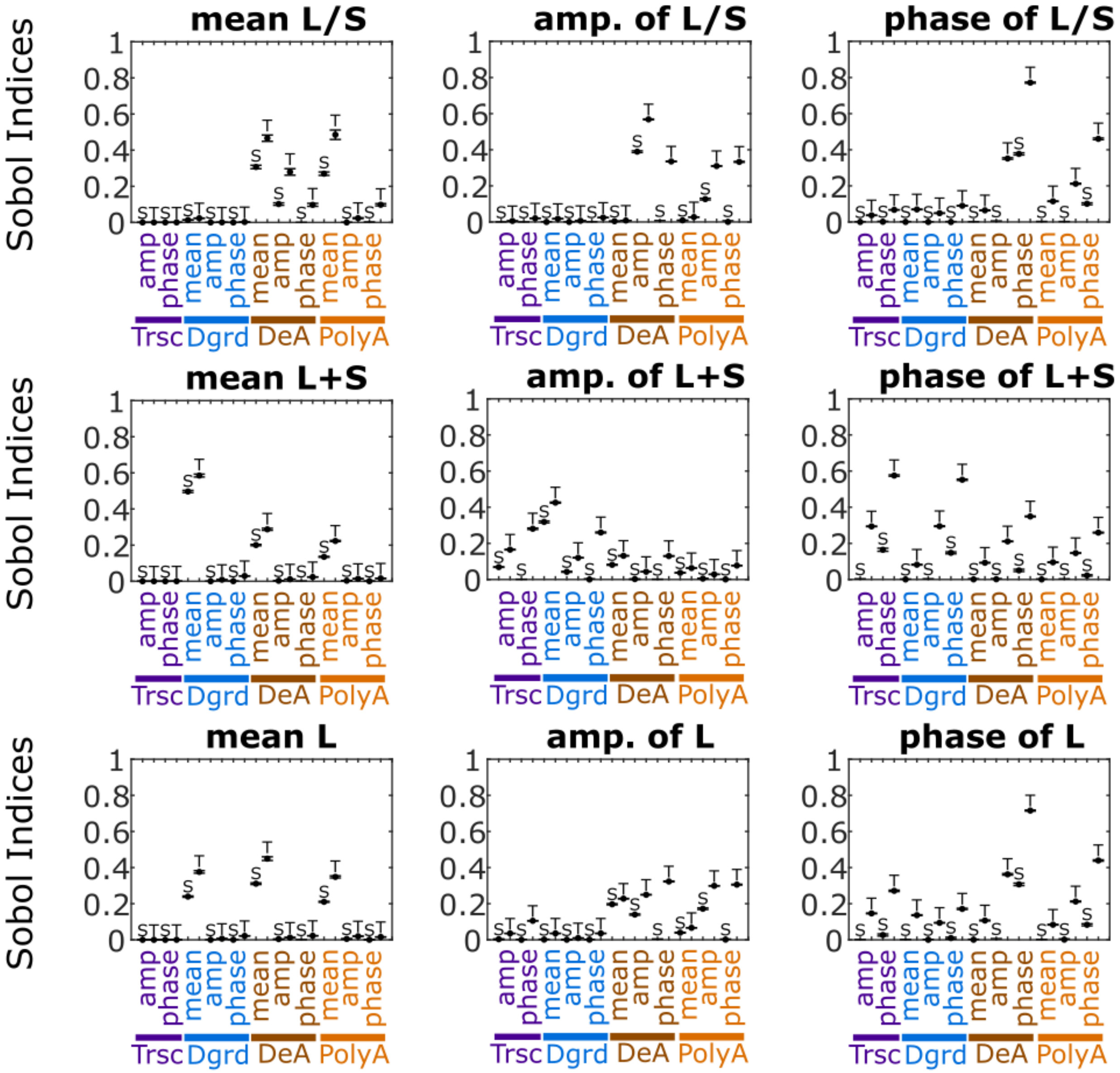


**Figure S2. Sobol indices of model with cytoplasmic polyadenylation.**

Label “S” on top: single Sobol indices. Label “T” on top: total Sobol indices. The mean (black dot) and standard deviation (black cap) of the Sobol indices from 10 repeats were plotted. Each repeat was performed using the procedure described in Methods with $N = 100{,}000$.

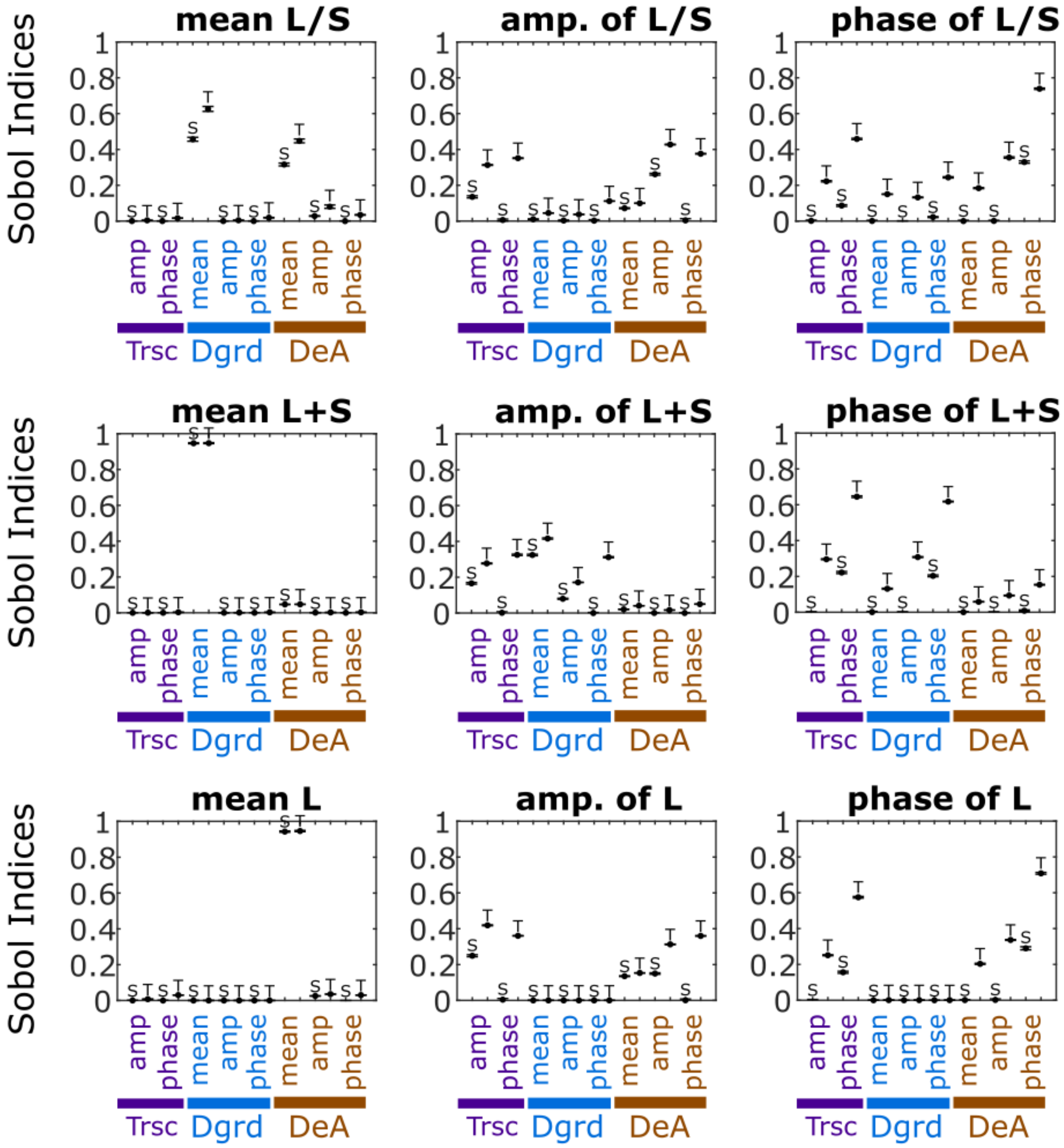


**Figure S3. Sobol indices of model without cytoplasmic polyadenylation.** Label “S” on top: single Sobol indices. Label “T” on top: total Sobol indices. The mean (black dot) and standard deviation (black cap) of the Sobol indices from 10 repeats were plotted. Each repeat was performed using the procedure described in Methods with $N = 100{,}000$.
